# Assessing changes in whole-brain structural connectivity in the unilateral 6-hydroxydopamine rat model of Parkinson’s Disease using diffusion imaging and tractography

**DOI:** 10.1101/2024.11.22.624742

**Authors:** Mikhail Moshchin, Roger J Schultz, Kevin P. Cheng, Susan Osting, James Koeper, Matthew Laluzerne, James K. Trevathan, Andrea Brzeczkowski, Cuong P. Luu, John-Paul J Yu, Richard Betzel, Wendell B. Lake, Samuel A. Hurley, Kip A. Ludwig, Aaron J. Suminski

**Affiliations:** Department of Neurological Surgery, University of Wisconsin-Madison, Madison, WI USA; Wisconsin Institute for Translational Neuroengineering, University of Wisconsin-Madison, Madison, WI USA; Department of Neurology, University of Wisconsin-Madison, Madison, WI USA; Department of Radiology, University of Wisconsin-Madison, Madison, WI USA; Department of Neuroscience, University of Minnesota, Minneapolis, MN USA

## Abstract

Parkinson’s disease (PD) is a multifactorial, progressive neurodegenerative disease that has a profound impact on those it afflicts. Its hallmark pathophysiology is characterized by degeneration of dopaminergic neurons in the midbrain which trigger a host of motor and non-motor symptoms. Many preclinical research efforts utilize unilateral lesion models to assess the neural mechanisms of PD and explore new therapeutic approaches because these models produce similar motor symptoms to those of PD patients. The goal of this work is to examine changes in brain structure resulting from a unilateral lesion both within the nigrostriatal system, where dopaminergic neurons are lost, and throughout the brain. Using multi-shell diffusion magnetic resonance imaging and correlational tractography, we assessed microstructural changes throughout the brain resulting from unilateral injection of 6-hydroxydopamine (6-OHDA) in the median forebrain bundle (MFB). Following lesioning, the PD phenotype was confirmed using behavioral and histological assessment. Correlational tractography found networks of fiber tracts that were either positively or negatively correlated with lesion status throughout the brain. Analyzing patterns of intra- and inter-hemispheric connectivity between the positively and negatively correlated fiber tracts revealed two separate neural networks. The first contained only negatively correlated fibers in the lesioned hemisphere consistent with the local effects of the lesion (i.e. dopaminergic depletion in the nigrostriatal system). The second contained systematically overlapping fiber tracts in the lesioned and non-lesioned hemispheres including the olfactory system and cerebellum, which we suggest are indicative of adaptive mechanisms to compensate for the lesion. Taken together, these results suggest that correlational tractography is a reasonable tool to examine whole brain structural changes in rodent models of neurodegenerative disease, and may have future translational value as a diagnostic tool for patients with PD.

## II. Introduction

By 2030, nearly 10 million people worldwide will be living with Parkinson’s Disease (PD) – a 66% rise driven by the aging global population. This progressive neurodegenerative disorder devastates lives through a cascade of events including the degeneration of dopamine producing neurons in the substantia nigra pars compacta (SNc) (1). As striatal dopamine deficiency worsens, patients face quality of life limiting motor symptoms including resting tremor, rigidity, slowness of movement and freezing of gait. Over the last two decades, studies have demonstrated that the non-motor effects of PD (i.e. olfactory dysfunction, gastrointestinal issues, sleep disturbances, depression, anxiety, pain, and fatigue (2) are equally troublesome and often manifest much earlier than the well-recognized motor symptoms leading to the assumption that the pathophysiology of PD has far reaching consequences throughout the brain. The use of advance imaging techniques has begun to address this question, showing that PD has significant effects on brain structure (3–6), function (7,8) and metabolism (9,10) beyond the basal ganglia networks typically associated with the disease. Unfortunately, these advances have not yet extended to preclinical models of PD where they would be beneficial for characterizing its progressive pathophysiology and the resulting dysregulation of neural circuits in the basal ganglia and throughout the brain.

Since its development in the 1960s (11,12), injection of the neurotoxin, 6-hydroxydopamine (6-OHDA), within nigrostriatal pathways (i.e. the median forebrain bundle (MFB), striatum or SNc) has been one of the most commonly employed preclinical models of PD. Here, oxidative stress caused by 6-OHDA serves to lesion nigrostriatal dopaminergic (DA) neurons (11) resulting in dopaminergic depletion and dysfunction similar to that observed in patients with PD. Importantly, rats with 6-OHDA lesions experience similar motor symptoms as those seen in patients with PD allowing for detailed investigations of the neuroanatomical (13), electrophysiological (14,15), and neurochemical effects (16) of PD as well as the validation of therapeutic interventions (17–19). Most commonly, only one hemisphere is lesioned with dopaminergic depletion evolving over approximately 14 days (13) and other systemic compensatory mechanisms over a periods of weeks to months (20–26). Despite the focal, asymmetric lesion of nigrostriatal DA neurons, some have reported broad, bilateral changes in both brain structure and function (27–31). However, many of these investigations are limited, in that the focus is on quantifying changes within spatially limited regions of interest (ROIs), neglecting changes in brain-wide networks associated with the lesion itself (i.e. dopamine depletion) and the resulting compensatory mechanisms.

To overcome the spatial constraints of earlier studies, recent advances imaging methods based on diffusion-weighted magnetic resonance imaging (dMRI) have opened new avenues for assessing the integrity of white matter (WM) tracts in diseased brain networks, and have proven to be valuable clinical tools for diagnosing various diseases (see (32,33) for reviews). The power of dMRI lies in its high sensitivity microstructural changes in the cellular environment. Measures of diffusion are often related to changes in the distribution of diffusing water molecules with increased anisotropy often being linked to highly organized structures like to large axonal pathways (34). These assumptions, however, are confounded by the presence of neurodegenerative disease and injury as measures of diffusion like fractional anisotropy (FA), mean diffusivity (MD), radial diffusivity (RD) and axial diffusivity (AD) are known to either increase or decrease based on the nature of the underlying pathology (see (32) for review). For example, in a study examining the relationship between measures of water diffusion and underlying histological changes in a model of traumatic brain injury, Chary, et. al (35) demonstrated that decreased in FA was directly related to properties of the injured tissue like decreased myelin density, loss of mylenated axons or increased cellularity (i.e. gliosis). Indeed, many studies have linked reduced mesures of anisotropy to a broad spectrum of diseases including PD (3,5,6,29,30,36–38), but increased anisotropy is less frequently reported and the underlying factors remain unclear. Some have linked increased anisotropy to adaptive or compensatory mechanisms like neuroplasticity (39), while others have suggested that it is related to greater number or density of myelinated fibers as is seen early in development (40,41) or disorders like PD (42–44), schitzophrenia (45), or Williams Syndrome (46).

This work directly quantifies structural changes resulting from unilateral dopaminergic depletion both local to the lesion site and throughout the brain. Here, we utilized ex-vivo, multi-shell, diffusion magnetic resonance imaging to assess microstructural changes in rats with unilateral lesions of the median forebrain bundle (MFB) triggered by a 6-OHDA injection. We performed behavioral tests and quantitative histology to evaluate the 6-OHDA induced Parkinsonian phenotype and utilized correlational tractography (i.e. connectometry)(47) to evaluate differences in brain structural connectivity between lesioned and non-lesioned rats. Our work shows that unilateral injection of 6-OHDA results in dramatically reduced diffusion (i.e. negative correlation with lesion status) local to the lesion, which is consistent with disruption of the nigrostriatal pathway. Importantly, brain microstructure changes were not limited to the site of injection, but rather extended bilaterally throughout the cerebral cortex, midbrain, and cerebellum with areas of both increased and decreased diffusion (i.e. positive and negative correlation with lesion status) observed. Based on analysis of intra- and inter-hemispheric patterns of connectivity, we suggest that structural changes resulting from unilateral lesions are not limited to specific nodes within the basal ganglia but rather extend throughout the brain and result from both direct degeneration of the nigrostriatal pathway and subsequent compensatory mechanisms. Our findings demonstrate that connectometry measures are appropriate for evaluating the effects of neurodegenerative disease in preclinical models, offering a promising tool for the future exploration of the neural mechanisms PD and therapeutic interventions in both preclinical models and human patients. A preliminary exploration of these data have appeared as a conference proceeding (48).

## III. Methods

### Experimental Design

A total of 12 male Long-Evans rats (12 weeks of age, weighing 313-403 g) were used in this study. Rats were housed on a 12-hour light/dark cycle with ad libitum access to food and water. The experimental timeline (Fig. 1A) consisted of 4 distinct phases designed to quantify behavioral, system, and cellular level neural changes resulting from unilateral dopaminergic depletion. At day 0, rats were randomly assigned into two groups, Vehicle (n = 6) and PD (n=6), and then underwent stereotactic surgery to inject either saline or 6-OHDA into their left medial forebrain bundle (MFB). Beginning 19 days post-surgery, motor performance of each rat was measured using standard behavioral techniques (Fig 1C) to quantify the effects of dopaminergic depletion (12,49). Following the completion of behavioral assessments, the rats were perfused and had their brains extracted for subsequent diffusion MRI (Fig 1D, Day 28) and histological examination of nigrostriatal neurons via tyrosine hydroxylase (TH+) staining (Fig. 1E). All animal procedures were reviewed and approved by the Institutional Animal Care and Use Committee (IACUC) at the University of Wisconsin-Madison. One rat in the Vehicle cohort was lost to a surgical complication, and one rat in the PD cohort was excluded from analysis as it did not show rotational behavior when challenged with APO. Thus, all analyses included data from n=5 rats in the Vehicle cohort and n=5 rats in the PD cohort.

**Fig. 1.**
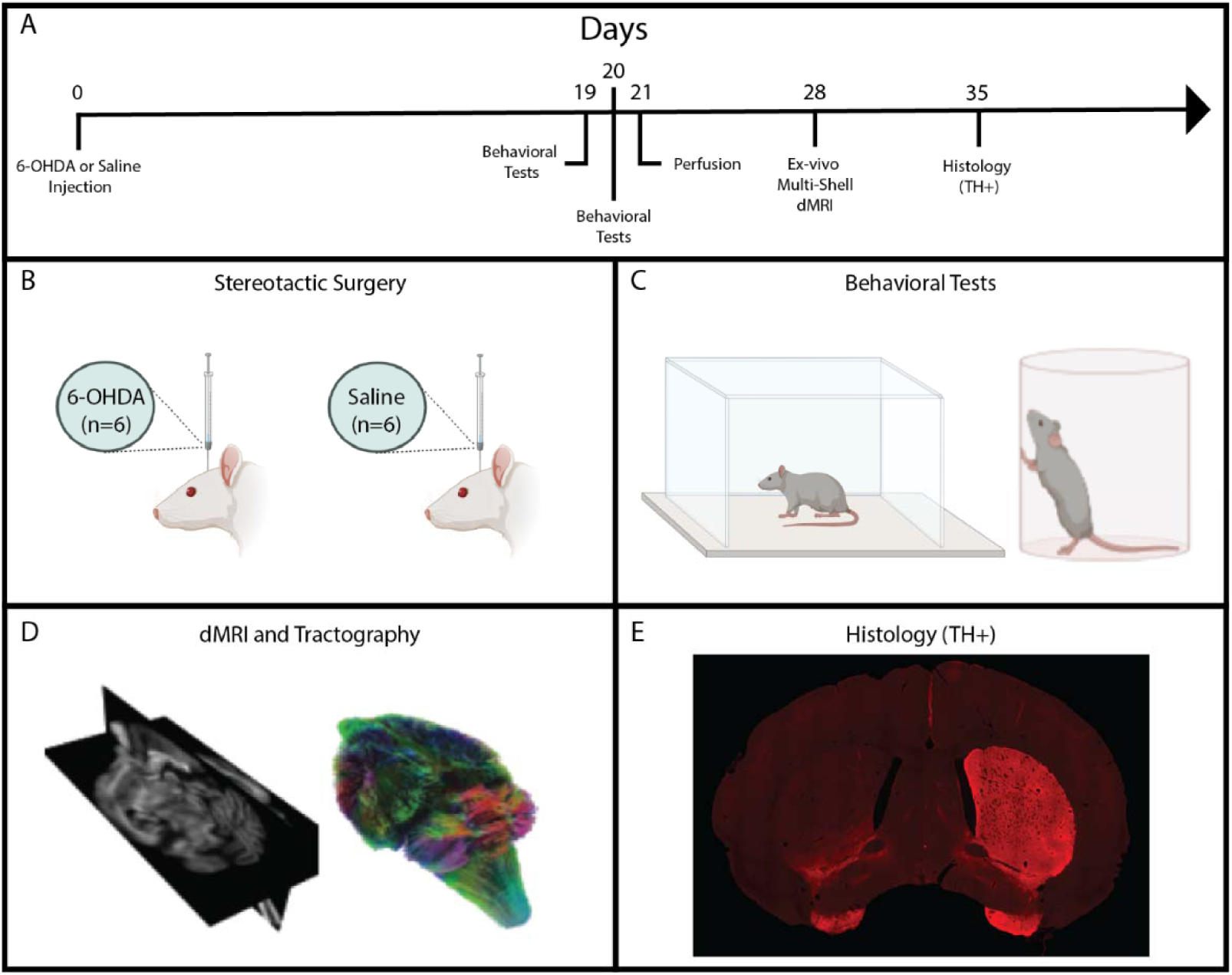
Experimental timeline and design. **A)** Timeline of the experimental procedures. **B)** On day 0, sterteotactic surgery was performed to inject saline in the Vehicle rats (n=6) and OHDA in the PD rats (n=6). **C)** On days 19 and 20, behavioral testing was performed to asses the motor function of all rats. Forelimb use asymmetry was assessed in a cylindrical field; testing for rotational locomotion following apopmorphine (APO) injection was done in an square open field. **D)** On day 28, ex-vivo, multi-shell dMRI was acquired. **E)** 35 days post-injection, brains were stained with tyrosine hydroxylase (TH+) to identify asymmetry in degeneration of striatal dopaminergic neurons.

### Surgical Procedures

Rats were anesthetized with isoflurane in O_2_ (induction at 5%, maintenance at 1-2.5%). Following induction, rats were placed in a stereotaxic frame, the top of the head was shaved, and the skin prepped with alternating povidone iodine and alcohol scrubs. At lease 30 minutes prior to stereotaxic inejctions, Desipramine HCL (20 mg/kg) was administered to prevent neurotoxic damage to noradrenergic neurons (50). The skin was incised along the midline and tissue was resected exposing lambda and bregma. After the skull was leveled, a drill was used to create a small burr hole above the injection location (-4.3mm AP, −1.6mm ML) (51). A microscope was used to visually confirm contact of the needle with the dural surface prior to the needle being lowered into the MFB. After reaching the desired depth (-8.0mm DV relative to the dura), the needle was left to rest for 5 minutes. Rats in the 6-OHDA cohort received injections of 6-OHDA (8 μg in 2 μL of saline with 0.02% ascorbate over 8 min), whereas rats in the vehicle cohort received injections of sterile phosphate buffered saline (2 μL over 8 min). After completing the injection, the needle was left to rest for 5 minutes before it was withdrawn. Gel foam soaked in sterile saline was used to cover craniotomy and 5-0 vicryl suture was used to close the skin incision. Throughout surgery, a heating pad was used to regulate body temperature, and the heart rate, SPO2, and body temperature of the rats were monitored (PhysioSuite, Kent Scientific).

### Behavioral Tests

Two behavioral assays were performed on days 19 and 20 to quantify motor function after stereotaxic injections, the forelimb use asymmetry/cylinder test, and the apomorphine induced rotational (APO) test. Both tests are routinely used to evaluate motor performance in dopamine depleted rats (49). To complete the cylinder test, rats were placed in a glass cylinder (160 mm in diameter and 395 mm in height) without prior acclimatization and videorecorded for 5 min. The number of times a rat used its forelimbs to navigate the wall were identified and counted. Forelimb contact with the wall was categorized as either ipsilateral to the lesion, contralateral to the lesion, or both. Touches were quantified as percentages and computed as: (ipsilateral touches)/(ipsilateral touches + contralateral touches + both)*100; (contralateral touches)/(ipsilateral touches + contralateral touches + both)*100; and both/(ipsilateral touches + contralateral touches +both)*100. Two-sample t-tests were used to assess differences in the percentage of ipsilateral, contralateral, and simultaneous touches between the Vehicle and PD groups. A bonferonni correction was used to control for multiple comparisons.

The apomorphine rotational test is a predictive test that confirms the PD phenotype in a rat mode (12,20,52). On days 19 or 20, the rats were subcutaneously injected with apomorphine (0.1 mg/kg dissolved in 0.2% ascorbic acid and prepared in 0.9% sterile saline), placed in an open plexiglass box (303x303x303 mm), and video recorded for 30 minutes (53,54). During this test, rats with a sufficiently lesioned MFB are expected to rotate more frequently and away from the lesioned side (contralateral rotations), whereas non-lesioned rats should rotate significantly less with no preferred direction. Distance traveled, average speed, and number of contralateral rotations were quantified using the automated motion tracking package, DeepLabCut (see Video Tracking Analysis for details). Similar to the cylinder test, a two-sample t-test was used to compare the number of rotations between the PD and Vehicle groups.

### Video Tracking Analysiss

DeepLabCut, an open-source motion tracking platform (55), was utilized to effectively track the movement of a rat within an open field arena. A total of 240 frames from 12 videos were used to train a neural network that underwent 500,000 iterations. From the perspective of the video camera, four easily identifiable body parts were marked and used to train the neural network: the tip of the nose, right ear, left ear, and the base of the tail. 95% of the data set was used for the training the neural network, while the remaining 5% was used for validation. Tracking error was estimated to be 2.34 pixels (1.85mm) for the training data and 3.3 pixels (2.6mm) for the validation data. For each marker, DeepLabCut provided an X, Y, and detection likelihood value. A Savitsky-Golay filter was used on the X and Y coordinates to smooth fluctuations in tracker position. If the detection likelyhood value was less than 95%, the data were censored and replaced with a linearly interpolated value between the most recent coordinate above the accepted detection value and the subsequent coordinate that was also above threshold. Smoothed estimates of the X and Y position of each marker were averaged. This averaged position was used in subsequent analyses. The total distance traveled was calculated by summing the difference in pixels of the center of the animal between every two frames and converting this pixel value to centimeters (1 pixel = 0.079cm). Speed was calculated by identifying the center of the rat in two consecutive frames and then dividing that by the frame duration. Contralateral rotations were calculated by creating a vector between the nose and base of the tail of the animal and calculating the angle of this vector with respect to the x-axis. Each time the animal completed a full 360° rotation from its initial position in the contralateral direction, the number of contralateral rotations would increase by one. Counts of contralateral rotations were binned each minute of the recording.

### Perfusion

Perfusions were completed on the 21^st^ day following surgery. Here, rats were anesthetized with isoflurane (2-5%) and transcardially perfused with ice-cold PBS, followed by 4% paraformaldehyde (PFA). Following perfusion, the brain was extracted and placed in 4% PFA for 48 hours and then washed and transferred to 1x PBS.

### Diffusion MRI - Image Acquisition and Preprocessing

On day 28, each brain was removed from 1x PBS and placed in a 10 ml syringe filled with Fluorinert (3M). Syringes were grouped in batches of three and then imaged using dMRI. Prior to imaging, the orientation of the syrniges within the scanner was documented photographically to preserve the identity of each sample within the grouped 3D volume. Multi-slice diffusion-weighted spin echo images were acquired with a 4.7 T Agilent MRI system and a 3.5cm diameter quadrature volume radiofrequency (RF) coil. The following scanning parameters were used: TR = 2500 ms, TE = 28 ms, matrix = 128x128 (reconstructed to 256 x 256), field of view (FOV) = 32x32 mm, voxel size = 250 microns, section thickness = 0.35 mm (resampled to 0.25 mm isotropic in post-processing), number of excitations (NEX) = 2, and two shell acquisitions with 10 directions at b = 0 s/mm², 25 directions at b = 816 s/mm², and 50 directions at b = 2,041 s/mm² (D =12.20 ms, d = 6 ms). All imaging was performed in a temperature-controlled room at a consistent 20-21 °C.

Imaging data were converted to the NIfTI format and headers were manually verifed to match acquisition parameters (0.25 mm, 0.25 mm, 0.35 mm). FSL eddy-correct was then applied (FSL, FMRIB Software Library; Version 5.0; Oxford Centre for Functional MRI of the Brain; http://fsl.fmrib.ox.ac.uk/fsl/) to correct for image distortions caused by eddy currents. The 3D volume was then imported into DSI-Studio, resampled to an isotropic (ISO) voxel size (0.25 mm in each direction), and the individual brains were then separated and saved for individual processing. Next, individual images were aligned to AP-PC orientation using the fslreorient2std tool and the FSLeyes nudge tool was employed to individually adjust the affine transformation of each image to align them to the standard template. The sum of all transforms was then applied using FSL FLIRT (FMRIB Linear Image Registration Tool) to resample the image into the standard atlas template space (56). B-table accuracy was verified using an automated tool (57). Finally, diffusion data were reconstructed and spin distribution functions (SDF) computed using a model-free methodology, generalized q-sampling imaging (GQI)(58). Diffusion sampling length ratio was set to 0.2. The reconstruction was performed using the WHS_SD_Rat Template built into DSI studio.

### Diffusion MRI – Analysis and Tractography

A primary interest was to quantify how dopaminergic depletion may change structural properties of neural circuits both local to the lesion and throughout the brain. First, we computed and extracted various voxel-wise diffusion measures within regions of interest (ROIs) in the basal ganglia, thalamus, and sensorimotor cortex. Diffusion measures included: quantitative anisotropy (QA), fractional anisotropy (FA), mean diffusivity (MD), axial diffusivity (AD), and radial diffusivity (RD). ROIs were derived from the atlas defined by Johnson, et. al. (56). Within each ROI, we compared diffusion measures across the PD and Vehicle groups using two-sample t-tests. Statistical significance was determined at the α = 0.05 level.

Next, we used correlational tractography to assess local changes in fiber tracts that have significant correlation with disease state. Here, local connectome matrices were estimated by sampling SDFs for each animal using the local fiber directions of the Johnson atlas. Local connectome matrices were assembled into a connectome database and associated with a cohort (i.e. PD and Vehicle) using a regression model (47). Here, the regression model took the form, Y=XB, where Y is the n-by-m local connectome matrix, n is the number of animals, m is the total number of fiber directions in the atlas, X is a n-by-2 matrix recording 6-OHDA status (1 for PD, and 0 for Vehicle) and a vector of 1’s to estimate the intercept, B is a 2-by-m coefficient matrix. Local connectomes with positive or negative associations were tracked with a deterministic fiber tracking algorithm (59) using default parameters in DSI-Studio (T-score threshold = 2.5). The resulting tracks were filtered using a combination of topology-informed pruning (4 iterations, see (60)) and a 10 voxel (2.5mm) length threshold. A false discovery rate (FDR) of 0.05 was used to select tracks with significant association to a cohort. FDR was computed in DSI-Studio using the method described by Yeh, et al., (47). A total of 4000 permutations to the row vectors of B were used to estimate the empirical and null track length histograms.

The correlational tractography analysis resulted in two separate statistical network maps describing fiber tracts that have significant positive and negative correlation with disease state (i.e. PD vs. Vehicle). Given that our PD model depleted dopamine unilaterally, we wanted to understand if correlated fibers spread bilaterally across the brain or were restricted to the lesioned hemisphere. Pattens of connectivity were assessed by computing the connection strength between each ROI within the Johnson atlas. Here, connection strength was defined as the total number of tracts that end in two ROIs divided by the median length of the tracts. We used the Pearson’s correlation coefficient and Mantel test (61) to quantify the intra- and inter- hemispheric spatial correlation between connectivity matrices derived from the connectometry analysis. A total of 100,000 permutations of the rows and columns of the connectivity matricies were used to compute the Mantel statistic, which measures the agreement/similarity between two matricies. Next, we examined if performance on behavioral assessments and histological metrics were predictive of changes in the network maps. We extracted the average QA values for each animal associated with the positively and negatively correlated fibers. We then fit linear regression models (fitlm in Matlab) to examine the relationship between QA and 1) the histological ratio of mean-pixel intensity in the striatum coronal slices, and 2) the percentage contralateral paw touches. Fits were considered significant if an F-test found the slope to be significantly different from zero at α = 0.05. A Bonferroni correction was applied to adjust p-values for multiple comparisons.

### Histology

Following dMRI, brain tissue was sectioned through the basal ganglia and stained for tyrosine hydroxylase (TH). Tissue was cryoprotected in a 0.1M phosphate buffer with 2% dimethyl sulfoxide (DMSO) and a graded series of glycerol concentrations at 4°C as follows: 10% for 1 day, 15% for 4 hours, and then 20% for 4 days. The tissue was frozen with dry ice and sectioned in the coronal plane at 40 μm thickness with a sliding microtome. The slices were then transferred to an ethylene glycol based cryoprotectant at -20°C where they were stored. Slices were first washed in 1X TBS. Antigen retrieval was performed at 80°C in Tris-EDTA (pH 9.0), followed by endogenous peroxidase quenching in TBS with 3% H2O2 and 10% Methanol. Sections were washed in TBS with 0.05% Triton X-100 (TBST; Sigma-Aldrich, Burlington, MA), blocked in TBST with 5% normal goat serum for 30 minutes, and then incubated overnight in TBST with 1% bovine serum albumin (BSA; Calbiochem by Millipore Sigma, St. Louis, MO) and primary antiserum (Millipore Sigma MAB318; 1:5,000). After the washes in TBST, sections were incubated in Alexa Fluor 594 secondary IgG (Invitrogen A11032; 1:750) for 2 hours. Afterwards, the slices were rinsed with TBS and mounted on a gelatin coated slide and then cover slipped with Prolong Gold antifade and DAPI (Invitrogen by Thermo Fisher Scientific, Waltham, MA).

Coronal slices were imaged using a Axio Imager Z2 microscope and ZEN 3.3 software (Carl Zeiss). Images were obtained with a TxRed filter cube and a shutter speed of 4 ms at 50% LED power. For quantification of striatal TH appearance, mean-pixel intensity of TH signal was analyzed using the software ImageJ (62). For each brain, we analyzed three coronal slices of the striatum (approximately 1.00 mm, 0.20 mm and −1.00 from Bregma along the AP axis). For each slice, we outlined the caudate putamen (CPu) in both hemispheres manually and extracted the mean pixel intensity of each ROI. Subsequently, we averaged the values for both hemispheres of each slice and compared the ipsilesional to contralesional intensity by dividing the average ipsilateral value by the average contralateral value.

## IV. Results

Depleting dopamine unilaterally via injection of 6-OHDA is a commonly used technique to create a rodent model of PD that likely has marked effects on neural circuits beyond those directly impacted by the 6-OHDA lesion itself. Our goal in these experiments was to characterize structural changes in fiber tracts throughout the brain that result from unilateral dopaminergic depletion.

### Verification of Parkinsonian Model

We first used a battery of behavioral and histological measures to verify creation of the Parkinsonian phenotype (model) in rats injected with 6-OHDA (See Table 1 for summary testing data from each rat). Using the cylinder test, we found a dramatic change in forelimb use between the PD and Vehicle cohorts (Figure 2A). When compared to the Vehicle cohort, rats in the PD cohort demonstrated significantly reduced use of their contralateral forelimb (T_8_ = 5.02, p = 0.001) as well as simultaneous use of both forelimbs (T_8_ = 3.255, p = 0.012) to explore the walls of the cylinder. Correspondingly, rats in the PD cohort also significantly increased use of their forelimb ipsilateral to the 6-OHDA lesion (T_8_ = −4.44, p=0.002).

**Fig. 2.**
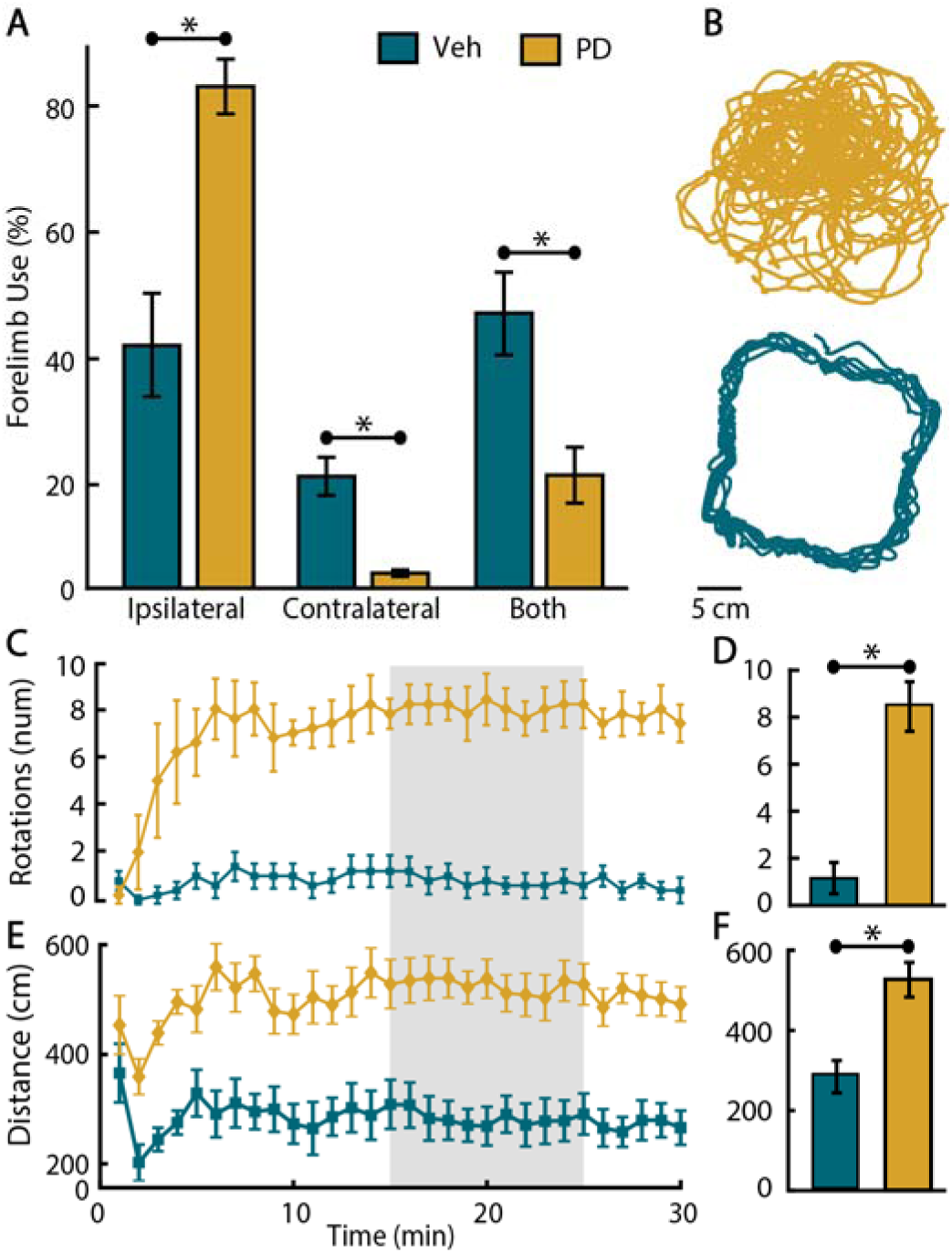
Evaluation of forelimb use and locomotor function in the PD and Vehicle cohorts. **A)** Forelimb use for rats the Vehicle (green) and PD (gold) cohorts. Contrasting percentage of touches that used the ipsilateral (*p=0.001), contralateral (*p=0.012), or both (*p=0.002) forelimb(s) demonstrate the significant asymmetry expected after unilateral lesion. **B)** Movement trajectories for Vehicle and PD cohorts after receiving a 0.1mg/kg apomorphine (APO) injection. Each curve represents the nose position of a rat, tracked with DeepLabCut. **C)** Average number of contralateral rotations for the Vehicle and PD cohorts in minutes 1 to 30 after APO injection. **D**) Average number of rotations for the Vehicle and PD cohorts in the 15 to 25 minute window (panel C, gray bar). Rats in the PD cohort rotated significantly more than those in the Vehicle cohort (*p<0.001). **E)** The mean distance traveled per minute over a 30- minute period after the APO injection. **F)** Average distance traveled for the Vehicle and PD cohorts in the 15 to 25 minute window (panel E, gray bar). PD rats traversed a significantly greater distance between minutes 15 and 25 than those in the Vehicle cohort (*p=0.004). All error bars represent ± 1 standard error of the mean.

**Table 1.** Summary behavioral and histological data for the Vehicle (V, n=5) and PD (n=5) groups.

| Group | Wall Touches |  |  | L/R Intensity Ratio (%) | APO Rotations (turn/min) |
| --- | --- | --- | --- | --- | --- |
|  | Ipsilateral (%) | Contralateral (%) | Both (%) |  |  |
| V1 | 64 | 7 | 29 | 91 | 3.1 |
| V2 | 47 | 24 | 29 | 77 | 0.4 |
| V3 | 25 | 18 | 57 | 76 | 0 |
| V4 | 39 | 17 | 44 | 84 | 1.4 |
| V5 | 18 | 23 | 59 | 101 | 0.2 |
| PD1 | 78 | 3 | 19 | 42 | 10.5 |
| PD2 | 88 | 4 | 8 | 32 | 7.2 |
| PD3 | 73 | 2 | 25 | 34 | 6.8 |
| PD4 | 68 | 2 | 30 | 47 | 6.4 |
| PD5 | 91 | 1 | 8 | 45 | 10.5 |

Examining locomotion following APO injection revealed characteristic rotational behavior commonly observed in dopamine depleted rats. Figure 2B shows representative movement trajectories for rats in the PD (gold traces) and Vehicle (green traces) groups. Notice that PD rats tend to walk in small circles in the middle of the open field (Fig 2B), whereas rats in the Vehicle group explore the perimeter of the arena. We computed the average number of rotations per minute for both groups and found that the number of contralateral rotations in PD rats began to increase 2 minutes following injection and reached a steady state approximately 5 minutes later. This rotational behavior continued throughout the entire 30-minute evaluation period. In contrast, rats in the Vehicle group exhibited minimal rotattion during locomotion. We computed the average number of rotations in the interval from 15-25 minutes (Fig. 2C, gray bar) after APO injection for an overall estimate of rotational behavior (Fig 2D). As expected, rats in the PD group demonstrated a significant increase in the rotational frequency compared to the Vehicle group (2 sample t-test, T_8_ = 6.76, p = 1.44e-04). We also examined the total distance traveled for all rats in each group following APO injection. Similar to the rotational measures, we computed the average distance traveled in each minute of the 30-minute testing period and summarized these data using the average distance traveled in a 10-minute interval beginning 15 minutes after APO injection (Fig. 2E and F). Again, rats in the PD group covered significantly more distance during the testing interval compared to the Vehicle group (2 sample T-test, T_8_ = 4.08, p = 0.0035).

In addition to the behavioral differences between cohorts, there were also stark changes in histological staining for dopamine in the striatum. Figure 3A shows representative stained slices for both groups. TH+ neurons are robustly labeled (Fig 3A, red fluorescence) in the bilateral striatum of rats in the Vehicle group whereas only the right striatum is labeled in the PD group. To quantify this asymmetry, we computed the ratio of left-to-right pixel intensities in the striatum for both the PD and Vehicle cohorts (see L/R Intensity Ratio in Table 1). A two sample T-test found a significant reduction of intensity ratio in PD rats compared to Vehicle (Fig 3B, T_8_ = 8.22, p = 3.59e-05) confirming that injection of 6-OHDA in the MFB destroyed many nigrostriatal dopaminergic neurons within the left hemisphere.

**Fig. 3.**
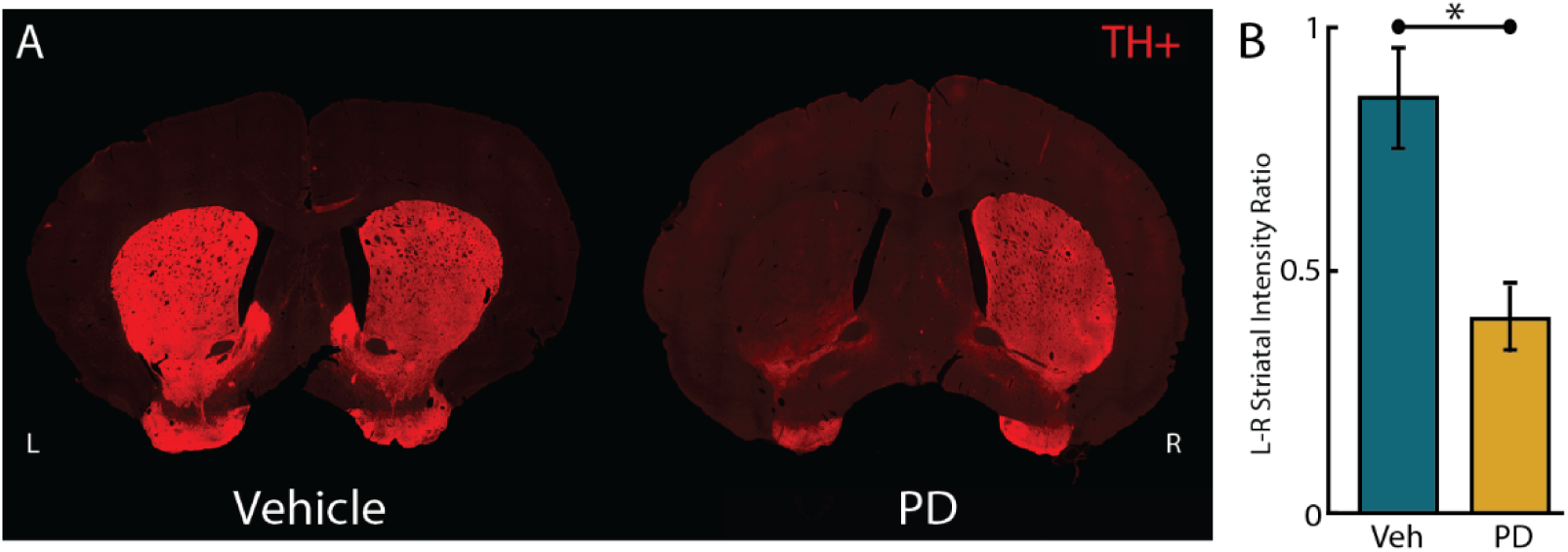
Histological examination of striatum coronal slices. **A)** Coronal slices of the striatum from a representative rat in each of the Vehicle and PD cohort. Bright red regions indicate neurons staining positive for TH. Notice the substantial reduction of DA neurons (i.e. TH+) in the left striatum of the PD sample. **B)** Ratio of mean-pixel intensity between the left and right striatum for the Vehicle (green) and PD (gold) groups (*p<0.001). Error bars represent +/- 1 standard error of the mean.

### Assessing Changes in Brain Microstructure due to Dopamine Depletion

After verifying that both behavioral and histological measures from the PD group are consistent with a high degree of dopaminergic depletion, we sought to characterize how neural circuits both local to the lesion and throughout the brain are impacted in this unilateral model of PD. Importantly, visual inspection of diffusion images between the left SNc and striatum, adjacent to the injection site, showed no obvious signs of damage in either the PD or vehicle group. Figure 4 shows representative coronal slices from GQI reconstructed diffusion images from each animal.

**Fig. 4.**
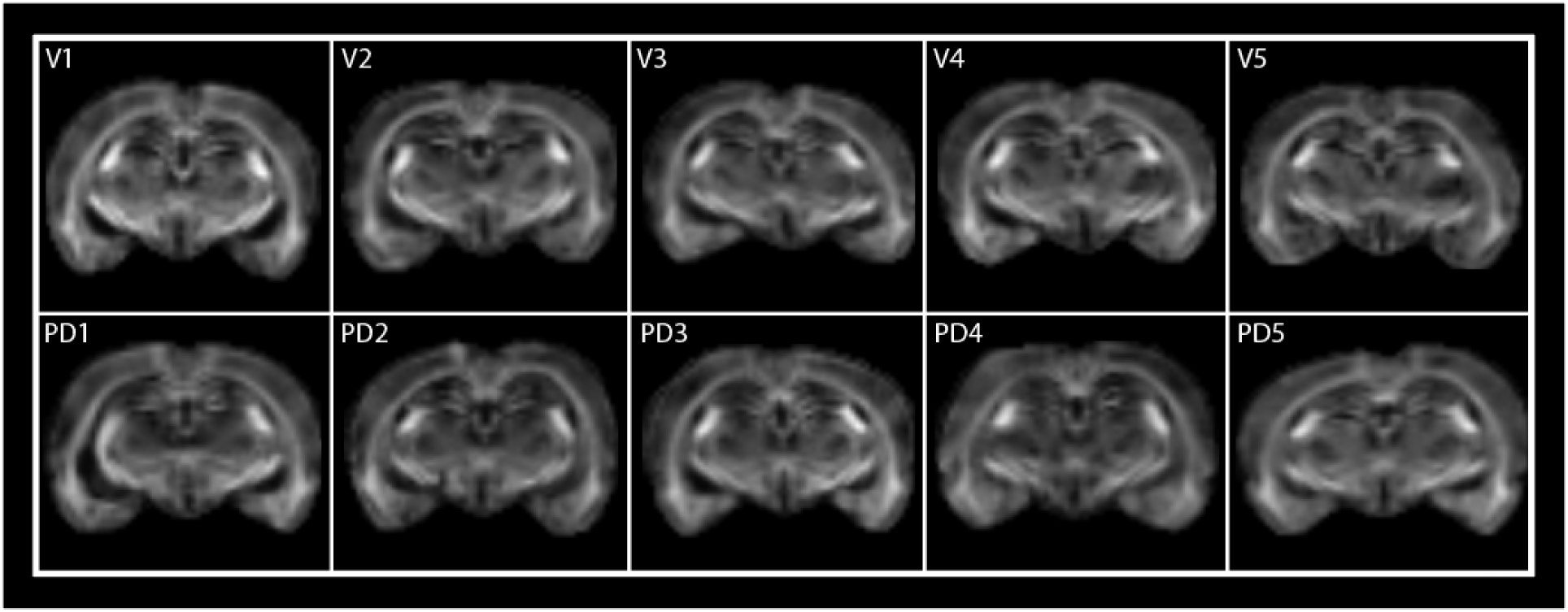
Representative coronal slices from each rat in the Vehicle (V1-V5) and PD(PD1 – PD5) cohorts.

We used multiple voxelwise and connectomic measures to assess changes in diffusion caused by unilateral depletion of dopamine via injection of 6-OHDA in the left MFB. First, we extracted multiple measures of diffusion from 13 distinct ROIs within the cortico-thalamo-basal ganglia (BG) network (see Table 2 for included regions). Two-sample t-tests were used to assess changes in each diffusion measure (QA, FA, MD, AD, RD) between the vehicle and PD groups.

**Table 2:**
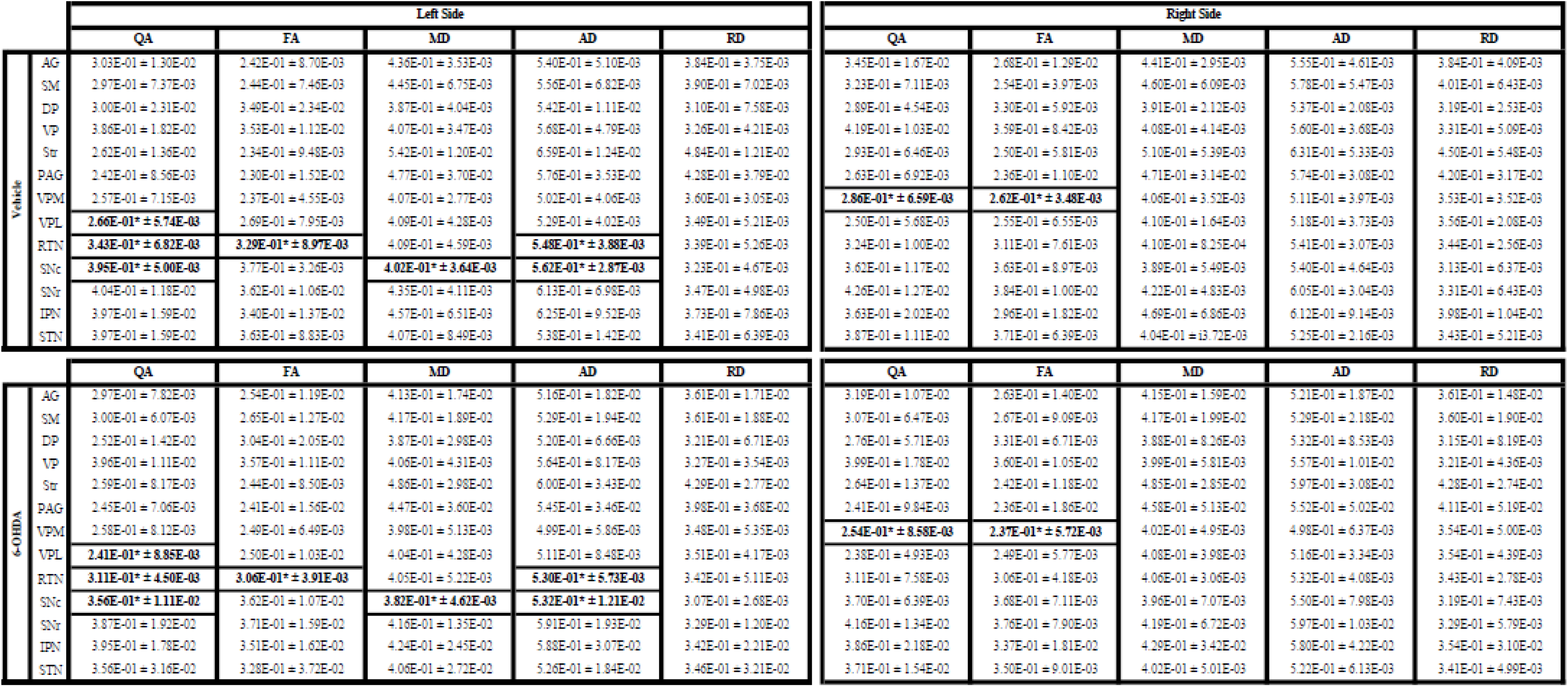
Voxel-wise diffusion measures within regions of interest (ROIs) in the basal ganglia, thalamus, and sensorimotor cortex.

The mean (± 1 standard error) values for all diffusion measures in each ROI is shown in Table 2. In the left hemisphere, uncorrected, two-sample t-tests found a significant decrease (p < 0.05) in mean QA of the PD group compared to Vehicle within the substantia nigra pars compacta (SNc), ventral posterior lateral nucleus of the thalamus (VPL), and thalamic reticular nucleus (RTN). Similar decreases were noted for MD and AD in the SNc and FA and AD in the RTN. Analysis of ROIs in the right hemisphere shows a significant decrease in QA and FA in the ventral posterior medial nucleus of the thalamus (VPM) for the PD group compared to Vehicle (p < 0.05). No significant differences in RD between PD and Vehicle were observed.

In addition to the voxelwise measures of diffusion, we locally sampled SDFs from the PD and Vehicle cohorts for connectometry analysis (47). Here, our goal was to estimate the effects of unilateral dopamine depletion throughout the brain by directly comparing local microstructural changes between the PD and Vehicle cohorts. Initial visual inspection of the resulting local connectome matrices (Fig. 5A, B) did not exhibit obvious differences between the two groups. We used correlational tractography to examine local changes in fiber tracks that were correlated with group membership (i.e. PD or Vehicle). Connectome length histograms were computed for both the empirical and null distributions for positively and negatively correlated fibers (Fig 5C and D). Differences in the distributions were used to estimate a fiber track length threshold (10 voxels) ensuring an FDR < 0.05 (Fig 5E).

**Fig. 5.**
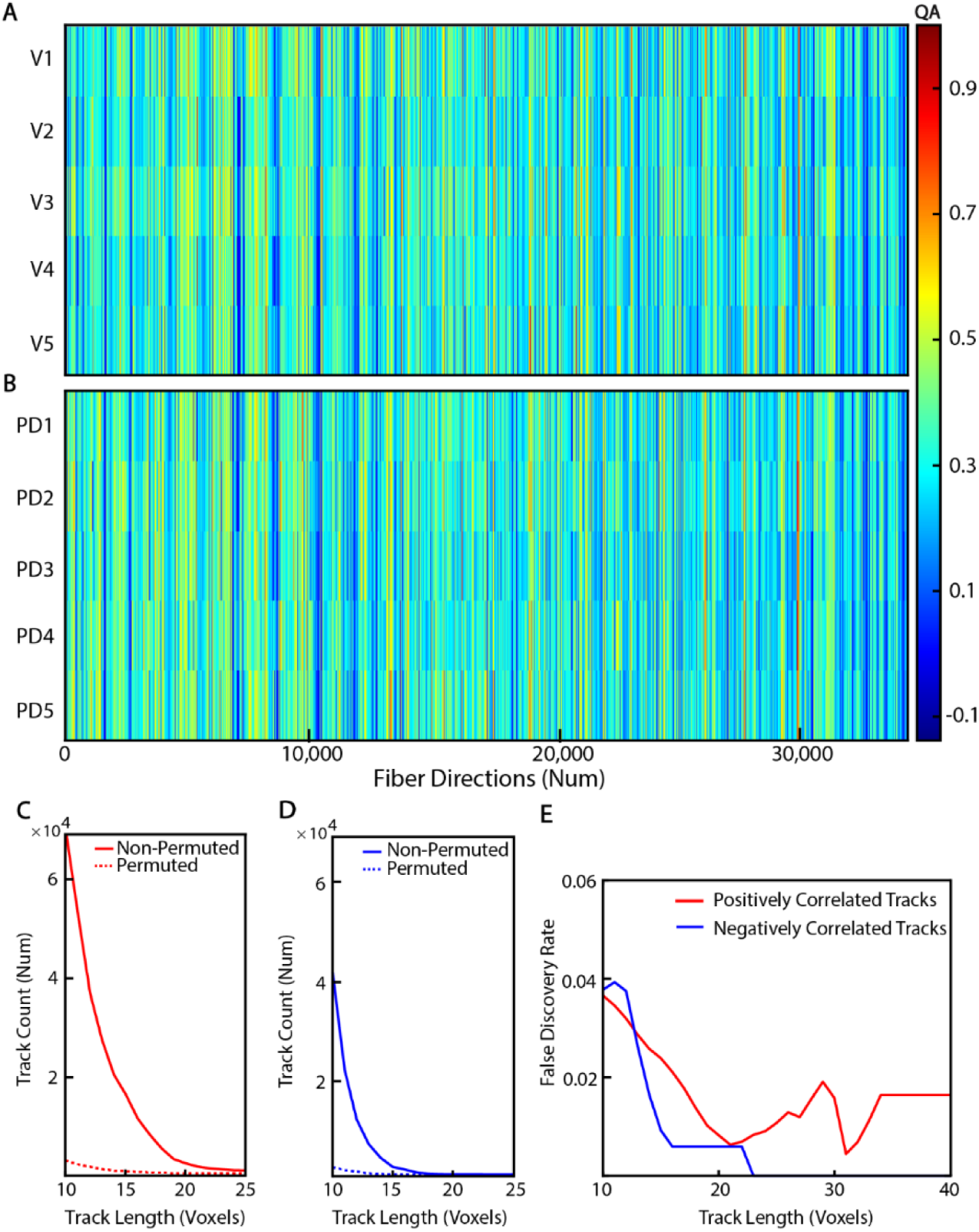
**A and B)** Visualization of vectorized local connectome matrices from 10 subjects. Each sample is characterized as a row vector of size 34332, where the number of row represents the number of local fiber directions in the reference atlas. **C)** Permutation tests examining the relationship between the number of positively (C) and negatively (D) associated fiber directions and track length were used to estimate the false discovery rate for the correlation measure (FDR < 0.05). The large difference between the track length histograms for the solid (non-permuted) and dashed (permuted) curves suggests that there are tracks associated with a substantial difference of the local connectome due to the 6-OHDA lesion. **E)** FDR remained less than the significance level (0.05) for track lengths longer than 10 voxels (2.5mm).

Correlational tractography found fiber tracks throughout the brain that both positively and negatively correlated with membership in the PD cohort (Fig 6A; red and blue fibers, respectively). Importantly, connectometry corroborated histological data showing significant unilateral dopamine depletion by demonstrating a negative correlation (i.e. reduced diffusion in PD rats compared to Vehicle) in the fibers connecting the left substantia nigra and striatum (Fig 6B; MFB, blue fibers). We also found negatively correlated fibers connecting the left striatum to the left ventral thalamus and fibers that crossed the midline to the non-lesioned hemisphere and reached the right ventral posterior medial nucleus of the thalamus (VPM). Two additional clusters of negatively correlated fibers were contained completely within the non-lesioned (right) hemisphere. The posterior cluster contained fibers near the right SNc that putatively connect the periaqueductal gray matter to the SNc and ventral thalamus. The anterior cluster connected the right striatum to widespread areas of the right frontal cortex, including the olfactory bulb, inferior frontal cortex (IFC: including the olfactory, gustatory, insular and piriform cortex), as well as the somatomotor and somatosensory areas. Like the negatively correlated fibers, correlational tractography found many fibers having a significant positive correlation (i.e. increased diffusion in PD rats compared to Vehicle) in both the lesioned and non-lesioned hemispheres. Most prominent were a large number of fibers throughout the cerebellar cortex, brain stem, and midbrain (Fig 6C, Cerebellum). The lesioned left hemisphere also showed substantial positively correlated fibers in large white matter tracts (corpus collosum, anterior commissure and internal capsule) and, of note, an area of the inferior frontal cortex that mirrored the negatively correlated fibers on the non-lesioned (right) hemisphere (Fig 6 B and C, IFC).

**Fig. 6:**
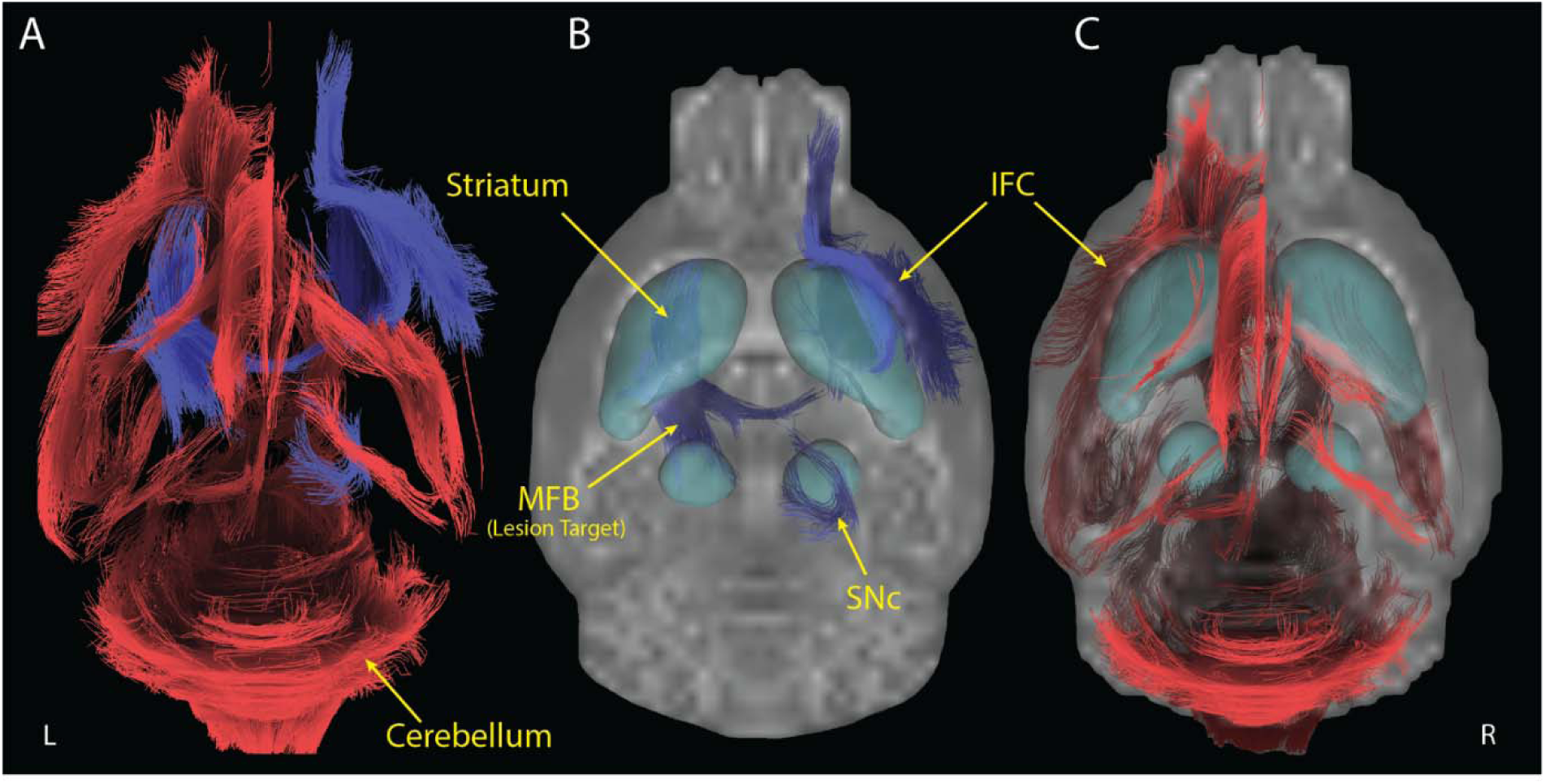
Correlational tractography describing changes in brain structure associated with unilateral 6-OHDA lesion of the left MFB FDR < 0.05). **A)** Overlayed network maps of fiber tracks positively (red) and negatively (blue) correlated with disease status (i.e PD or Vehicle). **B)** Negatively correlated tracks shown in isolation highlight changes in the ipsilesonal nigrostriatal pathway, including the median forebrain bundle (MFB) and contralesional inferior frontal cortex (IFC). **C)** Positively correlated tracks shown in isolation hightlight ipsilesional changes in the IFC and bilateral changes in large white matter pathwys, brainstem and cerebellum. Teal regions in B and C represent the striatum and substantia nigra.

Based on the apparent interhemispheric symmetry observed between positively and negatively correlated fibers in the lesioned and non-lesioned hemispheres (i.e. IFC and cerebellum), we sought to further understand if the microstructural changes caused by the unilateral 6-OHDA lesion involved similar regions in both hemispheres. To do this, we first estimated the region-to-region connectivity for the positively and negatively correlated fibers and then quantified similarity in the patterns of connectivity using a Mantel test. Connectivity matrices for the correlational tractography analysis are shown in Figure 7A with positively correlated connections (red dots) shown in the upper triangle and negatively correlated connections shown in the lower triangle (blue dots). Notably, both positively and negatively correlated fibers have a higher density of intrahemispheric connections with very few connections that cross the midline. Exceptions are mostly limited to positively correlated connections in large white matter tracts (Fig 7A, upper right quadrant).

**Fig. 7.**
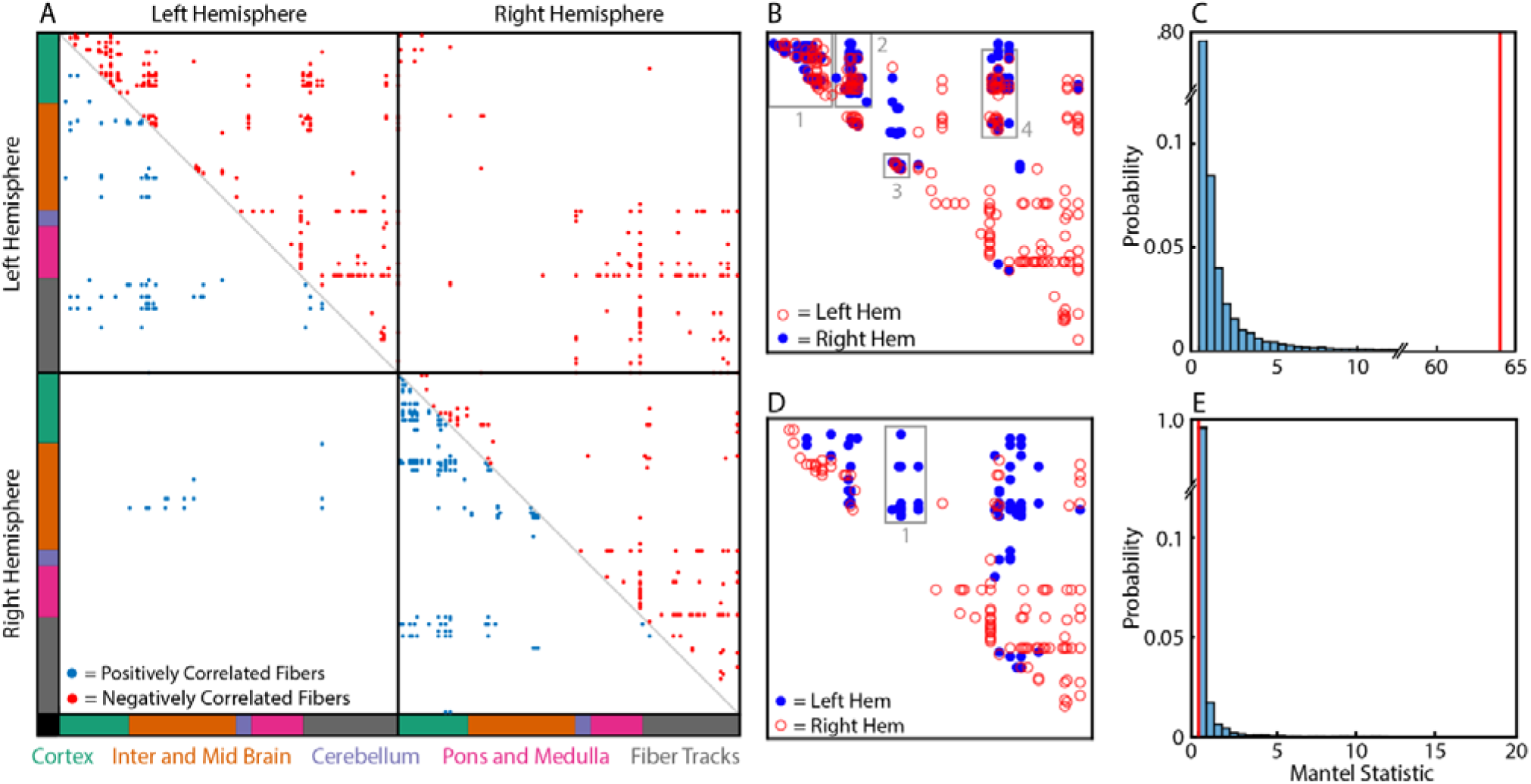
Connectivity matricies revel networks directly related to the lesion and subsequent compensatory changes. **A)** Positively correlated fibers(red) and negatively correlated fibers (blue) associated with the 6-OHDA cohort. **B)** The overlap between the lesioned (left) hemisphere positively correlated fibers and the non-lesioned (right) hemisphere negatively correlated fibers. Box 1 contained intracortical connections between the superior and inferior frontal cortex to the amygdala. Box 2 contained cortico-striatal connections and basal ganglia connections. Block 3 involved the substantia nigra connections. Block 4 contained large white matter tracts connections the frontal cortex, striatum, and ventral pallidum. **C)** The overlap between the distribution for the permuted Mantel statistic and the value non-permuted Mantel statistic (red line)demonstrates statistical significance (p=2.8*10^-4), meaning that there are systemic patterns of overlap between the connectivity matrices in B. **D)** The overlap between the lesioned (left) hemisphere negatively correlated fibers and the non-lesioned (right) hemisphere positively correlated fibers. Box 1 contained large white matter tracts connecting the left striatum and substantia nigra to other nuclei in the basal ganglia. **E)** The overlap between the distribution for the permuted Mantel statistic and the value non-permuted Mantel statistic (red line) demonstrates that there is no statistical significance (p=0.12), meaning that there are no systemic patterns of overlap between the connectivity matrices in D.

We began analyzing intrahemispheric connection patterns by comparing positively correlated fibers in the lesioned (left) hemisphere with negatively correlated fibers in the non-lesioned (right) hemisphere (Fig 7B). The Mantel test found significant systematic similarity in the patterns of connectivity (Mantel Statistic = 63.9, p = 2.1x10^-4^; see Fig 7C for empirical value and null distribution) driven primarily by overlapping connection in 4 regions (Fig 7B; Box 1 – 4). Box 1 contained intracortical connections between areas in the superior and inferior frontal cortex, including the cingulate, gustatory, olfactory, insular, somatomotor, somatosensory, orbital, and piriform cortex, and the amygdala (area labels from the Johnson Atlas; (56)). Box 2 contained cortico-striatal connections as well as those within the basal ganglia (striatum, nucleus accumbens and ventral pallidum). Connections within Block 3 involved the substantia nigra, while those in Block 4 connected large white matter tracts (anterior commissure, corpus collosum, internal capsule and olfactory tract) to the frontal cortex, striatum and ventral pallidum. We also observed significant similarity in the pattern of connectivity between positively correlated fibers in both the lesioned and non-lesioned hemisphere (Mantel Statistic = 682.5, p = 1x10^-5^) due to the large numbers of connections in the cerebellum and brainstem. Finally, we compared negatively correlated fibers in the lesioned (left) hemisphere with positively correlated fibers in the non-lesioned (right) hemisphere (Fig 7D). Unlike the previous comparisons, the Mantel test found no similarity between these patterns of connectivity (Mantel Statistic = 0.02, p = 0.197; see Fig 7E for empirical value and null distribution). Notably, the negatively correlated fibers, putatively related to the 6-OHDA injection (Fig 7D, Box 1; i.e. those connecting the left striatum and substantia nigra with large white matter tracts to other nuclei in the basal ganglia), were only seen in the lesioned hemisphere. Importantly, there was no change in the significance of the Mantel test when unthresholded connectivity matrices were used in computations.

As a final step, we performed a simple linear regression to evaluate the extent to which the dopaminergic depletion and limb use asymmetry could predict the quantitative anisotropy (QA) of fibers that significantly correlated with cohort. For both positively (Fig. 8A) and negatively correlated (Fig. 8B) fibers, regression found a significant relationship (i.e. slope) between QA and the ratio of left-to-right striatal, mean pixel intensities (T_9_ = −4.0, p = 0.004 and T_9_ = 2.8, p = 0.023, respectively). In contrast, only fibers positively correlated with cohort showed a statistically significant relationship between QA and the percentage of contralateral paw touches (Fig 8C; T_9_ = 2.8, p = 0.023). Negatively correlated fibers showed no no relationship between QA contralateral paw touches (Fig 8D; T_9_ = 1.53, p =.165).

**Fig. 8.**
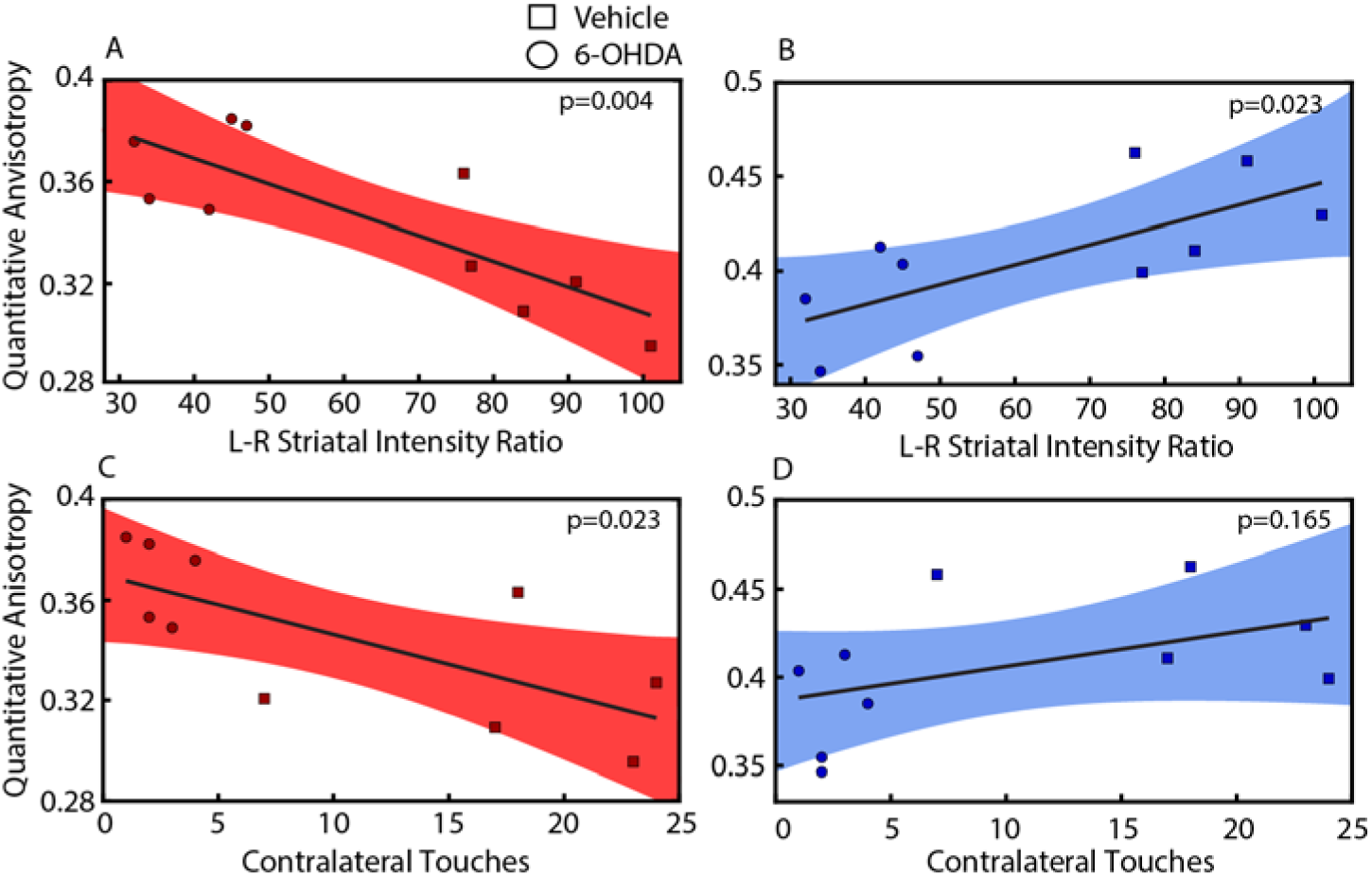
Regression of mean QA to intensity ratio or contralateral touches. **A-B)** Regression of QA from either positively (A) or negatively (B) correlated fibers to ratio of left-to-right striatal, mean pixel intensities after TH+ staining. **C-D)** Regression of QA from either positively (C) or negatively (D) correlated fibers to the number of wall touches by the forelimb contralateral to the MFB lesion. Solid lines represent a linear fit; red and blue region represent 95% confidence intervals.

## V. Discussion

While Parkinson’s Disease (PD) has traditionally been characterized as a motor disorder, manifesting visibly through resting tremor, bradykinesia, and gait disturbances (1). As such, both preclinical and clinical research has largely focused on quantifying changes in motor behavior and the neural control of movement due to PD and therapeutic interventions to treat it. Unfortunately, this focus has neglected the more recently documented non-motor effects of PD (63) and the fact that the pathophysiology of PD has far reaching consequences throughout the brain (64,65) far beyond the borders of the dopaminergic system. Our study aimed to examine structural changes resulting from PD, characterizing both the local sites of dopaminergic neurons lost within the nigrostriatal system and the broader brain regions where adaptive mechanisms likely manifested. Using the unilateral 6-OHDA rat model of PD, we employed multi-shell diffusion MRI and correlational tractography to compare whole-brain microstructural changes between lesioned (i.e. dopamine depleted) and non-lesioned rats. Importantly, we demonstrated that the effects of dopamine depletion (i.e. PD) extend beyond specific nodes (ROIs) within the basal ganglia, exhibiting both degeneration of the nigrostriatal pathway local to the lesion and subsequent compensatory mechanisms throughout the brain.

### Network Changes Reflect Nigrostriatal Dopamine Depletion

Depletion of dopaminergic neurons in the nigrostriatal system has far reaching effects on a broad repertoire of motor and cognitive functions, seen in both preclinical animal models (66) and humans with PD (63). In our model, unilateral dopamine depletion resulted in canonical motor behaviors described in previous reports. Correlational tractography found that behavioral changes were accompanied by many fibers negatively correlated with cohort (i.e. lower QA in PD group compared to Vehicle) connecting nuclei of the basal ganglia and thalamus in the ipsilesional hemisphere and the thalamus in the contralesional hemisphere. Reduced values of QA are typically associated with areas of neural injury/demyelination and have been shown to correlate with the clinical presentation of patients with neurodegenerative disorders (67). Most importantly, we found that the fibers of the MFB (Fig 6A), the site of 6-OHDA injection, were strongly negatively correlated with the PD group, a finding consistent with diminished TH+ staining in the ipsilesional striatum (Fig 3). Corroborating evidence between histological staining and imaging strongly support the ability of correlational tractography to measure changes in brain structure.

This is further supported by the fact that regression found a significant relationship between the mean QA of negatively correlated fibers in each animal and the degree of dopamine depletion in the striatum (Fig 8B). In addition, negatively correlated fibers extended outside of the lesioned nigrostriatal pathway and even into the contralesional thalamus. Cross-hemispheric projections from the nigrostriatal pathway have been described previously (68,69), as well as changes in cellular physiology and gene expression resulting from unilateral lesions (70–73). We suggest that negatively correlated fibers in the lesioned hemisphere are directly related to the lesion caused by 6-OHDA, and that correlational tractography techniques could be used to detect similar changes in patients with brain injury from PD. Future work should investigate if these results could be extended to less significant lesions similar to those associated with prodromal PD.

### … and Adaptive Mechanisms to Compensate for Dopamine Depletion

Beyond the primary impacts caused by 6-OHDA injection in the ipsilesional nigrostriatal pathway, we identified extensive microstructural changes thoughout cortical and subcortical regions of both hemispheres, showing either positive (i.e. higher QA in PD group compared to vehicle) or negative correlation with PD. Analysis of the patterns of connectivity resulting from correlational tractograpy revealed two especially interesting brain networks modulated by unilateral dopamine depletion. Here, a permutation test found that the interhemispheric patterns of connectivity had significant overlapping regions compared to randomized networks (Mantel test, Fig 7). The first notable overlapping pattern involved opposing changes in the olfactory network: positively correlated fibers in the ipsilesional hemisphere and negatively correlated fibers in the contralesional hemisphere, where fibers were found to connect the striatum to the inferior frontal cortex andolfactory areas (including the olfactory bulb) (Fig 7B, Box 1). Bilateral modulation of the olfactory network is interesting since it aligns with the hallmark presentation of olfactory dysfunction in prodromal phase of idiopathic PD (74), often appearing years before motor symptoms are detected This push-pull phenomena runs in parallel with previous studies examining the olfactory system after unilateral 6-OHDA lesions (76,77). Similar to our findings, the authors described increases in the number of dopaminergic neurons within the ipsilesional olfactory bulb compared to the contralesional bulb and control consistent with compensatory responses for disruptions in neurogenesis resulting from altered dopaminergic innervation of the subventricualar zone. The second strong overlapping patterns of connectivity involved positively correlated fibers in the cerebellum and brainstem of both hemispheres. While the role of the cerebellum in PD remains a topic of significant debate (see (64,65) for review consensus evidencesuggests that cerebellar changes may represent an indirect, potentially adaptive effect of dopamine depletion (78,79). Several functional imaging studies report overreactive participation of the cerebellum in PD patients (80,81), and others have connected these changes to adaptive mechanisms supporting the dysfunctional basal ganglia (82,83). In our data, it is unclear to what extent these adaptive mechanisms are due to the pathophysiology of PD or unilateral dopamine depletion. Further, our data demonstrates a potential confound in unilateral lesion models of PD. Specifically, that physiological changes are not limited to the lesioned nigrostriatal system or even to the lesioned hemisphere, but rather involve systemic changes in both hemispheres. This finding potentially calls into question the practice of using the contralesional hemisphere as a within animal control.

### Unilateral 6-OHDA Model and Diffusion Imaging

The 6-OHDA lesion model has been used extensively to examine the mechanisms of PD (12) and evaluate novel therapeutics (18,84) due to its resemblance to the cellular pathophysiology and behavioral phenotype of humans with PD (see (85) for review). In particular, unilateral dopamine depletion via injection of 6-OHDA along the nigrostriatal pathway results in characteristic motor behavior and histological changes reflecting the extreme loss of DA neurons in the ipsilesional striatum and SNc (> 95% and 80%, respectively,(49,86,87). Our behavioral and histological observations align with these reports: lesioned rats exhibited significantly increased contralesional rotations when challenged with apomorphine, marked limb use asymmetry and substantial reduction in TH+ neurons in the ipsilesional striatum (Figs 2 and 3). Though we did not directly quantify cell loss in the SNc, it is reasonable to expect significant degradation due to the lesion based on the striatal histology and impaired motor function (49). Given that, 1) the behavioral and cellular/molecular changes caused by 6-OHDA injection are known to be very stable at least 30 days post lesion (20) and that 2) brain tissue in our study was fixed during this stable period (i.e. 21 days post lesion), we can expect that our implementation of the unilateral 6-OHDA model is consistent with published literature (12,17,84,87–90) and representative of late-stage idiopathic PD (91).

Despite remarkable similarities in the motor behavioral phenotype and nigrostriatal histological changes, preclinical studies employing diffusion imaging techniques to quantify circuit level changes resulting from unilateral 6-OHDA lesions have reported varied and inconsistent results (29,92,93). This has been especially true for studies demonstrating changes throughout the of the nigrostriatal pathway (i.e. from SN to the striatum). For example, Soria et al., described a decrease in ipsilesional FA and AD in the substantia nigra, as well as bilateral increases in RD within the striatum (29). Similarly, Camp and Naja, showed that FA decreased in the ipsilesional substantia nigra after 6-OHDA injection (92). In contrast, Perlberg et al., reported bilateral alterations in motor cortex, globus pallidus, and striatum but not the substantia nigra by means of structural and functional MRI (30) . Our ROI analysis shows similar variability. We found decreased QA (but not FA) in the ipsilesional substantia nigra of lesioned rats compared to non-lesioned rats, however, these changes were also accompanied by additional decreases in various thalamic nuclei bilaterally. Notably, changes were variable based on the measure of diffusion, highlighting the inherent difficulties in examining changes between the PD and vehicle groups within predefined ROIs, and underscoring the need for alternative methodologies to estimate structural changes related to dopamine depletion. These considerations motivated our use of diffusion weighted imaging and correlational tractography to resolve the inconsistencies between the ROI analysis of different diffusion measures (i.e. FA, MD, etc). This methodology associates changes in diffusion within fragments of white matter pathways with an experimental variable (i.e. PD vs. Vehicle in this work) rather than in large ROIs defined a priori (47,94), thus allowing us to estimate local microstructural changes resulting from the experimental manipulations.

### Limitations and Future Directions

Using a novel tool, correlational tractography, we demonstrated that structural changes resulting from unilateral 6-OHDA injection in the MFB are not contained within nigrostriatal pathway but rather extend throughout the brain and reflect mechanisms of injury and subsequent compensation. There are several limitations regarding our experimental design and techniques that should be considered when interpreting the results of this study. First, it is an exploratory study with low numbers of animals in each group and limited statistical power. As a result, some of the comparisons made were potentially underpowered and thus used uncorrected p-values when multiple comparison correction might be more appropriate. Importantly, this caveat does not include our main behavioral (Fig 2), histological (Fig 3) and correlational tractography (Fig 5-7) findings which are robust despite the small sample size. Future confirmatory studies are necessary to validate these findings. We also acknowledge that local and brain-wide changes resulting from dopamine depletion are not solely related to brain structure, and include changes in electrophysiology, cell biology and gene expression (14,15,70–73). It will be important to understand how these changes are related to those of the brain’s microstructural changes we describe in this report. Finally, these data show that tractography may have value as a diagnostic tool to differentiate disease state as well as to explore the neural mechanisms of PD. Future work should evaluate its ability to ascertain the efficacy of therapeutic interventions in animal models to better understand how it may benefit the diagnosis and treatment of individuals suffering from PD.

## VI. Acknowledgements

Support for this research was provided by the University of Wisconsin – Madison Office of the Vice Chancellor for Research and Graduate Education with funding from the Wisconsin Alumni Research Foundation and the Department of Neurological Surgery.

KAL is a co-founder and equity holder for Neuronoff, Inc. KAL is also a co-founder and equity holder of NeuraWorx. KAL is a scientific board member and has stock interests in NeuroOne Medical Inc. KAL is also a paid member of the scientific advisory board of Abbott and Presidio Medical, and a paid consultant for the Alfred Mann Foundation, ONWARD and Restora Medical. All other authors have no conficts to declare.

## VII. Author Contributions

AJS, MM, KPC, WBL and JKT conceived of the experiment. MM, KPC and JKT performed the surgical procedures. MM, JKT, ML and AB collected the behavioral data. SO performed the histological procedures. SAH and JPJY developed the MRI acquisition protocol. MM, AJS, RJS, RB and JK analyzed and interpreted the data. MM, AJS, RJS, CPL, SAH, KL and SO drafted the manuscript. AJS and WBL secured funding for the project. All authors contributed to refining the manuscript and approved the final version.

